# A multiplexed and quantitative assay for small-molecule detection via photo-crosslinking of structure switching aptamers

**DOI:** 10.1101/2024.04.24.590963

**Authors:** Sharon S. Newman, Brandon Wilson, Liwei Zheng, Michael Eisenstein, Tom Soh

## Abstract

There is an unmet need for molecular detection assays that enable the multiplexed, quantification of smallmolecule analytes. We present xPlex, an assay that combines aptamer switches with ultraviolet-crosslinkable complementary strands to record target-binding events. When the aptamer’s small-molecule target is present, the crosslinkable strand is displaced, enabling PCR amplification and detection of the relevant aptamer. In the absence of that target, the aptamer is readily cross-linked to the strand, preventing amplification from happening. The resulting aptamer-specific amplicons can be detected and quantified in a multiplexed fashion using high-throughput sequencing. We demonstrate quantitative performance for a pair of small-molecule analytes, dopamine and glucose, and show that this assay retains good specificity with mixtures of the two molecules at various concentrations. We further show that xPlex can effectively evaluate the specificity of cross-reactive aptamers to a range of different small-molecule analytes. We believe the xPlex assay format could offer a useful strategy for achieving multiplexed analysis of small-molecule targets in a variety of scenarios.

## Introduction

Molecular quantification assays based on binding between target molecules and affinity reagents, such as the enzyme-linked immunosorbent assay (ELISA), are an essential tool for both clinical diagnostics and basic research. Several techniques enable highly-multiplexed detection of protein analytes, such as the proximity ligation assay (PLA) and proximity extension assay (PEA)^1,2^. These methods use DNA-labeled antibodies to generate analyte-specific barcode readouts that can then be measured by conventional amplification and high-throughput sequencing methods, enabling users to investigate hundreds of protein analytes in a single assay. No such analog exists for small-molecule analytes, however, because these methods rely on multi-epitope binding. Small molecules by definition exhibit limited numbers of epitopes, and simultaneous binding with bulky antibody reagents is generally not feasible. Aptamers are single-stranded nucleic acid-based affinity reagents that can recognize small-molecule targets with high sensitivity and selectivity, and could offer a powerful alternative to antibodies in this context. However, most efforts to date at developing multiplexed aptamer-based assays have relied on readouts such as fluorescence, enzymatic reactions, and electrochemical signaling, which are relatively limited in terms of how much they can be scaled up^3,4^. Consequently, there is an unmet need for generalizable methods that enable sensitive, multiplexable quantification of small molecules.

As a potential solution, we report the xPlex assay, which employs structure-switching aptamers^5^ to achieve sequencing-based quantification of multiple small-molecule analytes in a single sample. xPlex combines target-specific aptamer switches with chemically-modified complementary DNA strands that can be crosslinked to the aptamer by ultraviolet (UV) irradiation in the absence of target binding. Target-binding events induce a conformational change in the aptamer and displace the complementary strand prior to crosslinking, and can then be subjected to PCR amplification, whereas crosslinked aptamers cannot. This means that PCR amplification can generate a readout of the extent of target binding from multiple aptamers in parallel, and the quantitative signals from each analyte can subsequently be read out and analyzed by high-throughput sequencing (HTS). As proof of concept, we used xPlex to simultaneously quantify dopamine and glucose with high accuracy. We subsequently demonstrated the generalizability of xPlex by using this assay to assess the specificity of four aptamers previously generated by our group against molecules in the kynurenine metabolic pathway. We propose that this approach could provide the foundation for highly-multiplexed assays for quantifying diverse small-molecule analytes.

## Results and Discussion

### Assay development and validation

The xPlex assay is based on the selective production of a DNA reporter upon target binding to a structure-switching aptamer, where the subsequent readout can then be quantified via HTS (**Fig. 1a**). Such aptamers are designed such that in the absence of their target, they assume an unfolded structure due to hybridization of the aptamer stem to a complementary DNA strand that is present at excess concentrations (**Fig. 1b**, left). In the presence of target, the aptamer refolds into an analyte-binding conformation that displaces the complementary strand (**Fig. 1b**, right), and the extent of this conformation switch within a population of aptamers is proportional to the analyte concentration. We modify the complementary strand with a crosslinking moiety (CNV-K, **Fig. 1c**),^6,7^ such that exposure to UV-A (365 nm, 10 s at 4°C) forms a covalent bond between aptamer-hybridized strands. These complexes cannot be amplified by PCR because the crosslinked strand blocks access to the primer-binding site on the aptamer (**Fig. 1b**, bottom). The complementary strand cannot act as a primer itself due to an amino modification of the 3’ end. This means that PCR can only amplify those aptamers that have bound their target, and the resulting amplicons can be quantified with HTS in a highly parallelized fashion based on previously-defined calibration curves for each pair.

**Figure 1.**
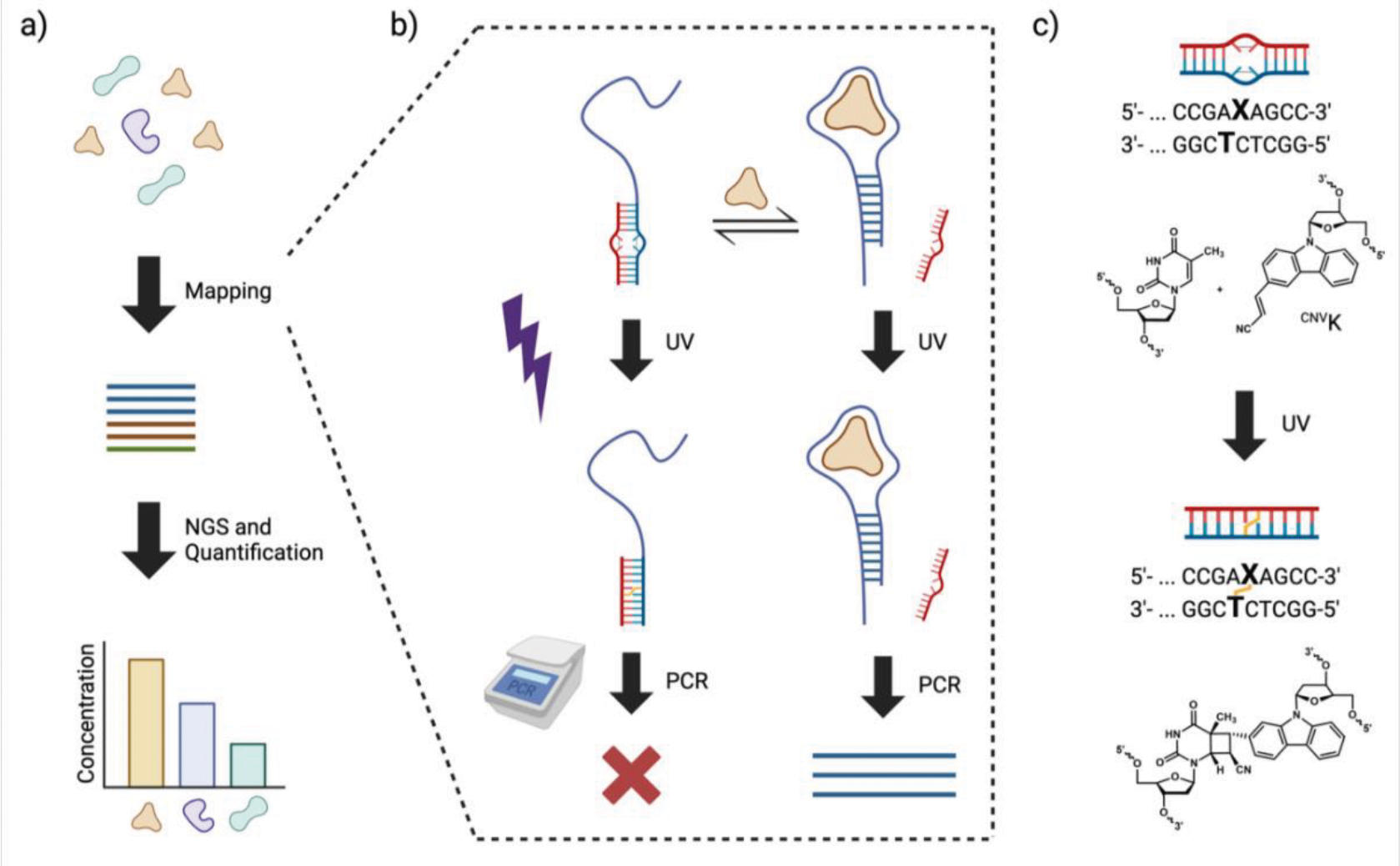
Overview of xPlex. **a)** xPlex produces a DNA reporter of small molecule-aptamer binding events, which can subsequently be sequenced and quantified. **b)** In the absence of target, structure-switching aptamers (blue) are bound to a competitor strand (red) modified with a ^CNV^K moiety (top left). Aptamer binding to its target (tan) displaces the competitor strand (top right). UV exposure covalently crosslinks non-target-bound aptamers to the competitor strand (bottom left). Only target-bound, non-crosslinked aptamers can be amplified with PCR (bottom right). **c)** Details of the crosslinking reaction; top sequence is the aptamer and bottom sequence is the competitor strand. Created with BioRender.

As an initial test, we used a previously published structure-switching dopamine aptamer^8^. We modified the competitor strand by inserting the CNV-K moiety near the 3’ end to reduce partial displacement by primers during PCR, and amino-modified the 3’ end to prevent the strand from acting as a primer itself (**Table S1**). After incubation with an excess of competitor (100 nM aptamer and 5 µM competitor), the aptamer ran at the expected size (65 nt) on a denaturing urea gel (**Fig. 2a**). After UV exposure, a high-molecular-weight band appears, indicating successful crosslinkage. We observed that a growing proportion of aptamers remained non-crosslinked in the presence of increasing concentrations of dopamine. We also observed a higher molecular weight band (∼80 nt) at higher dopamine concentrations that we attributed to crosslinkage of the aptamer to two competitor strands, as predicted by NUPACK^9^ secondary structure prediction at 4 °C (one at the hairpin stem, and another in the middle of the aptamer).

**Figure 2.**
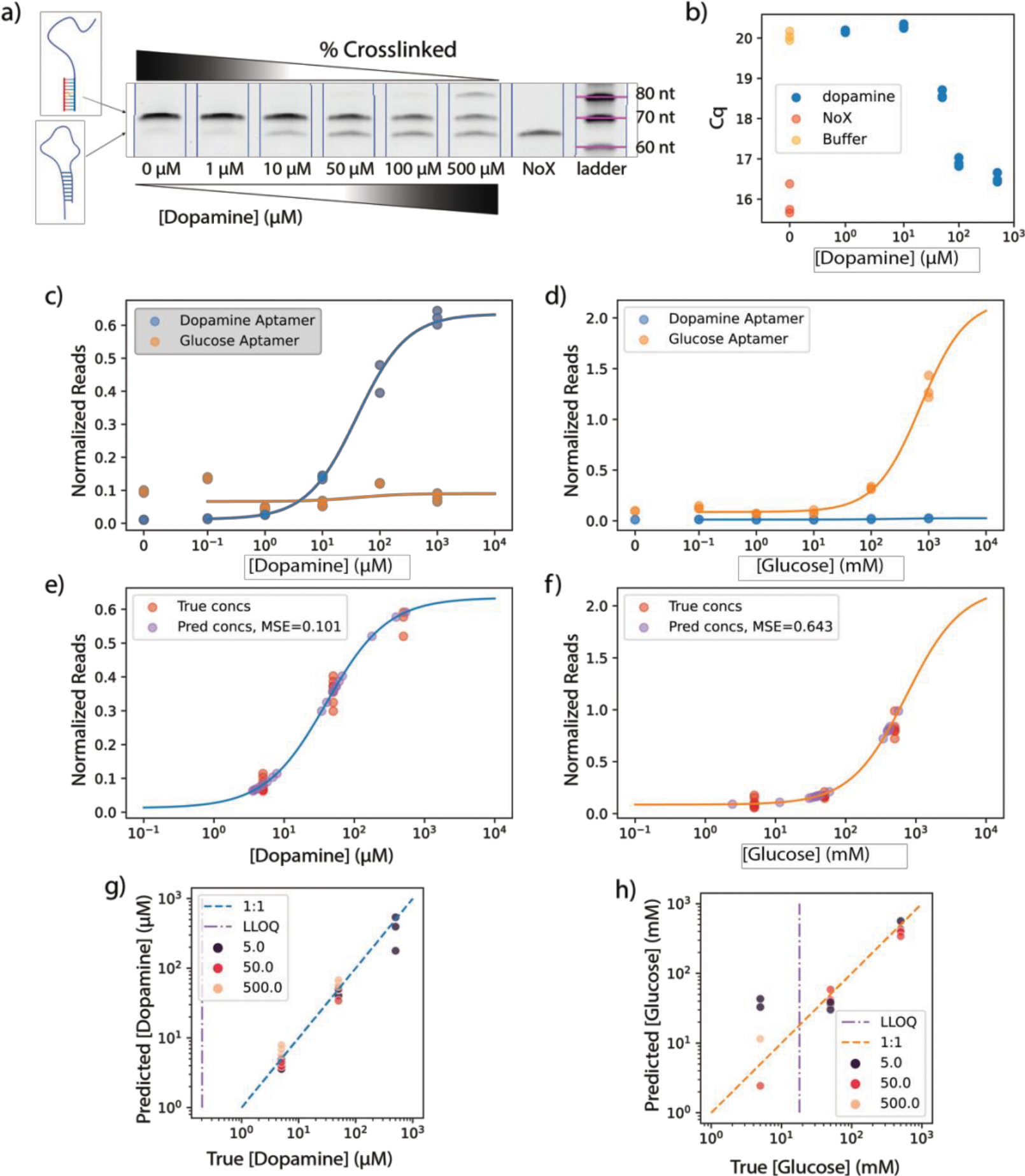
xPlex enables quantification of dopamine and glucose. **a)** Crosslinked aptamer concentrations decrease with increasing dopamine concentration, as seen on denaturing urea gels. Full gel is shown in **Figure S1**. NoX denotes uncrosslinked control. **b)** qPCR Cq values (n = 3) decrease with increasing dopamine concentrations. Dopamine-free controls with and without crosslinking are indicated in yellow (Buffer) and orange (NoX), respectively. **c, d**) Binding curves for dopamine and glucose aptamers in response to **c)** dopamine and **d)** glucose. **e, f)** Multiplexed quantification of **e)** dopamine and **f)** glucose in mixtures of these analytes at varying concentrations, with respective mean squared log errors (MSLE) based on 4PL fits for each aptamer. **g, h)** Plot of true versus calculated concentrations for **g)** dopamine and **h)** glucose. Dashed diagonal line shows 1:1 correlation. Vertical purple lines indicate lower limit of quantification (LLOQ). Color scale indicates analyte concentration of the non-target analyte (glucose in mM, dopamine in µM). Dots are individual replicates (n = 3).

We used qPCR to quantitatively confirm that crosslinking prevents amplification in a target concentration-dependent manner. Since the starting aptamer concentration is too high for qPCR, we first diluted the crosslinked samples 1,000-fold. Control samples of non-crosslinked aptamer and displacement strand showed a high starting DNA concentration (Cq = 15.9 cycles), whereas crosslinking of these two strands in the absence of dopamine yielded a higher Cq value (20 cycles), confirming that crosslinking greatly reduces the proportion of aptamer that can be successfully amplified (**Fig. 2b**). This difference in cycle number (while accounting for qPCR efficiency; **Fig. S2**) indicates that 95% of amplification is blocked by crosslinking. Increasing dopamine concentrations led to a commensurate decrease in Cq values, confirming that qPCR can provide a readout of dopamine concentration. However, given that gel electrophoresis and qPCR generates noisy data and are relatively low-throughput methods, we subsequently analyzed xPlex data via HTS.

We used custom primers (**Table S1**, RP1 and FP1) to add Nextera indexing tags so that multiple samples could be processed in one sequencing run. While converting our samples for sequencing, we can also introduce target multiplexing. We introduced a glucose aptamer^8^ into the sample mixture alongside the dopamine aptamer to demonstrate such multiplexing capabilities (**Table S1)**. Both aptamers are designed to displace the same competitor strand sequence when bound to their target, but since both aptamers have unique overall sequences, we can mix them together in equimolar amounts (with an excess of universal competitor strand) to simultaneously measure aptamer response with HTS. To minimize errors originating from PCR bias, we also introduced a control DNA sequence of known concentration into the PCR master mix, such that the raw reads for each indexed sample (**Fig. S3**) could be normalized to that control sequence. For quantitative assessment, we made separate serial dilutions for both dopamine and glucose and subjected these to binding, crosslinking, and HTS. For each indexed sample per analyte concentration, we had aptamer read-counts for both dopamine and glucose. Using the normalized sequence reads, we first fitted the dopamine aptamer reads in response to dopamine (**Figure 2c**) to a 4-parameter logistic (4-PL) curve, and calculated an effective equilibrium dissociation constant (*K*_*D*_) of 40 ± 6 μM (**Table S2**). This approximates the original reported *K*_*D*_ for this aptamer (10 µM), indicating that the addition of a crosslinking moiety and the xPlex process does not greatly affect its binding affinity. We also noted that the signal from our non-crosslinked control sample is greater than that from the aptamer subjected to the highest dopamine concentration in our assay. This means that the binding signal from that sample arises from actual aptamer saturation rather than a limitation of read depth, which is an important consideration for multiplexing (**Fig. S4)**. Finally, we determined that the lower limit of quantification (LLOQ) for dopamine in this assay was 200 nM, as determined by the 4-PL fitting parameters. For the glucose aptamer, we measured an effective *K*_*D*_ of 730 ± 1,637 mM for glucose, with an LLOQ of 18 mM (**Fig. 2d**). We note that the *K*_*D*_ for glucose is high with low confidence in the fit, which is due to the binding curve data not saturating. We simultaneously established the cross-reactivity of the aptamers by examining their response to non-target analytes. Both aptamers maintained consistent background levels despite increasing off-target concentrations (**Fig. 2c, d**), confirming the low cross-reactivity of these aptamers and demonstrating the feasibility of multiplexed analysis.

To demonstrate quantification of multiple targets within a single sample, we prepared seven different mixtures combining a range of concentrations of dopamine and glucose (**Table S3**). Based on the assumption of low cross-reactivity, the established binding curve fits, and reads of each aptamer within a sample, we were able to consistently calculate the original concentration of both targets (**Fig. 2e, f**). It should be noted that the low-confidence estimate of *K*_*D*_ for the glucose aptamer was not an impediment to analytical accuracy in these experiments. To assess our quantitative accuracy, we determined the mean-squared-log-errors (MSLE) of the calculated to true concentration values, which can loosely be interpreted as one minus the ratio of true versus calculated concentrations. We calculated MSLEs of 0.1 and 0.6 for dopamine and glucose. Importantly, the error in quantification of each analyte does not appear to be affected by the concentration of the other analyte (**Fig. 2g, h)**. We note that the lowest glucose concentration tested (5 mM) was below the LLOQ for glucose, and thus could not be accurately quantified (samples left of the purple line in **Fig. 2h**). Thus, although glucose detection was constrained by the poor target affinity of this aptamer, we confirmed that xPlex is capable of multiplexed quantification of small-molecule analytes.

### xPlex enables multiplexed reagent characterization

Finally, we examined how xPlex can be used to characterize the specificity of both target-specific and cross-reactive aptamers for small molecules within the kynurenine metabolic pathway. We employed four structure-switching aptamers previously obtained by our group^10^. Aptamers 3HK-1, XA-1, and KA-1 are selective for 3-hydroxykynurenine (3hk), xanthurenic acid (xa), kynurenic acid (ka), respectively. The fourth aptamer, SK-1, cross-reactively binds all three targets as well as kynurenine (kyn). The competitor sequence is universal to all four aptamers, and thus the only necessary step to incorporate them into xPlex was to add the CNV-K moiety to the competitor strand. To further optimize performance, we screened various competitor strand lengths and concentrations to maximize the signal-to-noise ratio (**Fig. S5–S8**). We established binding curves for the four aptamers and analytes using the optimized displacement strand concentration (500 nM) and length (17 nt). We measured effective K_D_ values of 100, 48, and 325 µM for 3HK-1/3hk, XA-1/xa, and KA-1/ka, respectively (**Fig. 3**). These aptamers exhibited high specificity, with no appreciable increase in signal from off-target analytes across a range of concentrations (**Table S4**). Our control assay without crosslinking confirmed that the saturated signals from our binding curves were from aptamer saturation and not a limitation of read depth, and showed minimal background noise (**Fig. S9**). The cross-reactive affinity reagent SK-1 exhibited an effective K_D_ of 149, 131, 40, and 260 µM for 3hk, xa, ka, and kyn, respectively. All reported K_D_ values are roughly an order of magnitude greater than previously reported values when these were also similarly fitted to a 4-PL curve (**Table S5**)^10^. This increase is a result of increasing the competitor strand length to lower the background signal. Nevertheless, the relative affinities of each aptamer for target and non-target analytes were similar to those measured by conventional assays, indicating that xPlex could offer a good mechanism for rapidly evaluating aptamer specificity in a high-throughput fashion. Importantly, despite higher effective *K*_*D*_s, the LLOQs were comparable: 7, 3, and 13 µM for 3hk, xa, and ka respectively with xPlex versus 0.5, 6, and 3 µM with the originally-published fluorescence assay, suggesting that xPlex-based quantification at lower concentrations should be minimally affected relative to each aptamer’s innate properties.

**Figure 3.**
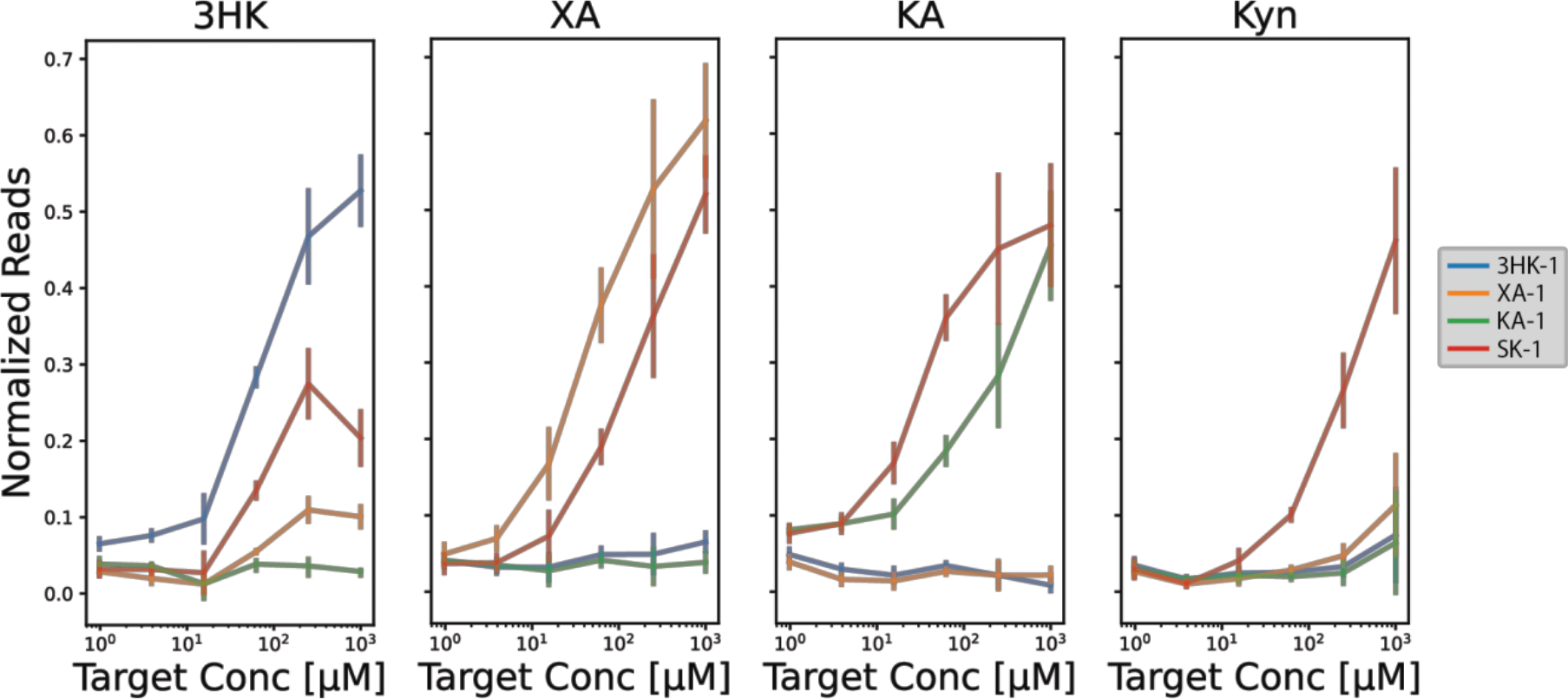
Characterization of kynurenine pathway aptamers with xPlex. HTS-derived binding curves for 3HK-1, XA-1, KA-1, and SK-1 with 3hk, xa, ka, and kyn. Each analyte concentration produced four separate reads for each aptamer, and all samples were processed in the same sequencing run. Target-free and no-crosslinking controls are shown in **Figure S9**. Non-normalized reads are shown in **Figure S10**.

## Conclusion

In this work, we introduce xPlex, a quantitative multiplexable assay for small molecules that uses structure-switching aptamers and crosslinking to generate a DNA readout of target binding that can be interpreted with conventional sequencing methods. We showed that we can quantify mixtures of dopamine and glucose with high selectivity and quantitative accuracy, while largely preserving the sensitivity of the original aptamers. Finally, we used four aptamers against various small-molecule components of the kynurenine metabolite pathway, and demonstrate that xPlex has potential as a high-throughput strategy for evaluating aptamer specificity for small-molecule analytes. Indeed, the selectivity and binding curve characterizations shown here would conventionally require separate analyses for each aptamer, whereas here we could simultaneously obtain four aptamer binding curves with only one serial dilution measurement. By producing unique DNA reporters representing aptamer and target binding fractions, our method could greatly facilitate multiplexed small-molecule quantification and aptamer characterization, where the only firm limitation is the sequencing read-depth and desired quantitative resolution.

## Methods

### Reagents

Aptamers, primers, PrimeTime probes, and adaptor sequencing strands (**SI Table S1**) were synthesized by Integrated DNA Technologies (IDT). Competitor strands were custom made in-house using 3-cyanovinylcarbazole phosphoramidite (CNVK), which was purchased from Glen Research. Kynurenine metabolites (L-kynurenine, XA, KA, 3HK), dopamine, and glucose were all ordered from Sigma-Aldrich.

PBS (10X), MgCl_2_ (1 M), Tween 20, Tris-HCL (1 M, pH 7.5), Axygen AxyPrep Mag PCR Clean-up Kits (MAGPCRCL), GelStar Nucleic Acid Gel Stain (#50535), and molecular-biology-grade 200-proof ethanol were purchased from Thermo Fisher Scientific. 2X GoTaq G2 Hot Start Colorless Master Mix was purchased from Promega (9IM743). Nextera XT Index Kits (96 indexes, 384 samples) were purchased from Illumina (FC-131-1002). SYBR Green I nucleic acid gel stain (10,000X concentrate in DMSO) (S7563) and Qubit dsDNA HS Assay Kit (Q32851) were purchased from Invitrogen. UltraPure TBE buffer (10X), Novex 10% TBE-Urea (EC68752BOX), 6x loading dye (R0611), and 20/100 single stranded DNA Ladder were purchased from IDT (#51-05-15-02).

### Instrumentation

Standard automated oligonucleotide solid-phase synthesis was performed on an Expedite 8900 synthesizer from Biolytic Lab Performance. DNA annealing and PCR reactions were done with an Eppendorf Mastercycler X50 96-well thermocycler. Crosslinking was done with a 365-nm Ultra High-Power Deep UV Curing Spot Lamp from Agiltron (SUVA-011). Gel images were captured with a ChemiDoc MP System from Bio-Rad Laboratories. Beads were prepared for HTS with DynaMag-96 side magnets purchased from Thermo Fisher Scientific (12331D). Viaflo 96-channel pipette, reservoirs, and 300 μl pipette tips were purchased from Integra BioSciences (6432).

### Oligonucleotide synthesis

Oligonucleotides were synthesized on an Applied Biosystems Expedite 8900 nucleic acid synthesis system. Oligonucleotide synthesis reagents were purchased from Glen Research. Commercially available fast-deprotecting phosphoramidites (Glen Research) were used for DNA synthesis of oligonucleotides containing CNVK modification. A 10 min coupling time was used for CNVK phosphoramidite. Deprotection of synthesized oligonucleotides was performed with concentrated aqueous ammonia for 2 h at room temperature, followed by concentration under reduced pressure. Oligonucleotides were purified using 20% denaturing PAGE and then eluted using the crush and soak technique. Finally, the eluent was desalted using C18-Sep-Pak cartridges and dried with a centrifugal vacuum concentrator. The 3’ end of the competitor strand was amino modified to prevent extraneous amplification.

### Binding and crosslinking protocol

For dopamine/glucose experiments, 100 nM aptamer was incubated with 5 µM competitor strand in 1x PBS and 2 mM MgCl_2_, unless otherwise indicated. The strands were pre-annealed by heating at 95ºC for five minutes in a thermocycler and dropping by 1ºC every 30 seconds until reaching 4 ºC. To obtain binding curves, we added 10 µl of pre-annealed competitor and aptamer to 10 µl of buffer and analyte, such that the final analyte concentration was in the range of 0–0.5 mM for dopamine, 0–500 mM for glucose, and 0–1 mM for the kynurenine analytes. For controls, we used equivalent volumes of buffer only. Samples were incubated on ice or at 4 ºC on a rotator for an hour. To crosslink, 10 µl of each sample was then transferred to a clear flat-cap 0.2-ml PCR tube and exposed to UV for 10 seconds, with the head of the lamp directly on the cap of the tube.

Aptamers used for dopamine and glucose data are ‘Comp15_dop_glu’, ‘Dop’, and ‘Glu’ (**SI Table S1**). Aptamers for kynurenine data are ‘3HK-1’, ‘XA-1’, ‘KA-1’, and ‘SK-1’ with ‘comp16’ unless otherwise indicated (**SI Table S1**). Kynurenine samples were prepared in buffer comprising 20 mM Tris-HCl, 120 mM NaCl, 5 mM KCl, 1 mM MgCl2, 1 mM CaCl2, and 0.01% Tween-20 in nuclease-free water. Aptamer and competitor were combined at 100 nM aptamer and 250 nM unless otherwise indicated.

To prepare samples for HTS, the protocol was modified as follows to increase throughput: 35 µl of pre-annealed aptamer-competitor mix was added to wells in a 96-well semi-skirted, LoBind twin.tec PCR plate (Eppendorf) containing 35 µl of analyte in the same buffer, such that each well achieved the appropriate range of analyte concentrations needed for binding curve generation in a final volume of 70 µl. The plate was sealed and incubated at 4 ºC on a rotator for 1 hour, and then spun down at 1,000 rcf for 30 seconds. Samples were then individually crosslinked for 10 s on ice with the head of the lamp directly on the surface of the seal, with a opaque paper covering samples that were not immediately being crosslinked. Since aptamer concentrations do not need to be as high for HTS, we used pre-annealed concentrations of 500 pM aptamer and 500 nM competitor.

### Gel electrophoresis

To prepare samples for electrophoresis, 4 µl samples were mixed with 4 µl of loading dye, then denatured at 95 °C for three minutes. The samples were run on 10% TBE-Urea Gels at 180 V for 50 min, stained with GelStar for five minutes, and then imaged.

### qPCR protocol

Samples were prepared as described above, then diluted 1,000-fold with buffer to avoid saturating the reaction. The qPCR mix was prepared with 1x GoTaq Master mix and 1x PrimeTime_dop probes which included RP1 and FP1(**SI Table S1**). 9 µl of this qPCR mix was mixed with 1 µl of sample and amplified using the following protocol: 95 ºC for 30 s, and 39 cycles of 95 ºC for 10 s, 51 ºC for 30 s, 72 ºC for 30 s, and then holding at 4ºC. We calculated the number of cycles required for each sample to reach 0.25 maximum fluorescent value (Cq). The amplification intensity value (*I*_*i*_) across all cycles was normalized as follows:

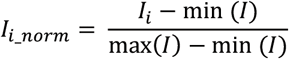

Using these normalized values, the cycle that first passes the threshold value of 0.25 is the Cq value. From the Cq values, we then calculated the fraction of unlinked to crosslinked samples,

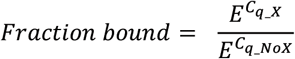

where *E* is the qPCR efficiency. To calculate *E*, we used PrimeTime probes (**Table S1**) to amplify known concentrations of dopamine aptamer and extract the Cq of each dopamine aptamer concentration. *E* is then calculated from the slope (m) of these Cq values across the log of the dopamine aptamer concentrations (**SI Fig. 2**):

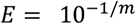

Based on this, we calculated E = 2.08. Finally, the percent of sequences that are blocked from amplification when crosslinked is:

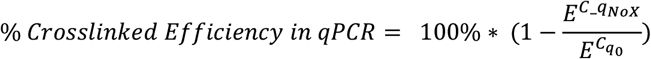

### Sequencing protocol

The sequencing protocol was modified from Newman *et al*.^11^ 45 µl of PCR reaction solution (1x GoTaq Master Mix, 500 nM forward primer, 500 nM reverse primer, and 15 pM control sequence) was added to each well in a fresh 96-well plate. For dopamine and glucose, we used FP1, RP1, and CTRL.15. For kynurenine analytes, we used FP17, RP2, and CTRL.17. The aptamers used for the kynurenine analytes were 3HK.17, XA.17, KA.17, and SK.17, with a 17-nt competitor strand (comp17) to provide additional stability. We used custom primers with adapters to apply Nextera Indexes later in the protocol; CTRL.17 is a custom control sequence with a random 15-nt region for post-HTS normalization with molecular barcoding (**SI Table S1**).

Samples with analytes, aptamers, and competitor were incubated and crosslinked as earlier described; however, all aptamers were mixed in the same solution at the stated final concentrations. 5 µl of sample was then added to each well. The plate was vortexed for one minute and spun down for one minute at 1,000 rcf and placed in a Mastercycler X50. Preamplification consisted of holding at 95 ºC for 30 s, and then 4 cycles of 95 ºC for 10 s, 51 ºC for 30 s, and 72 ºC for 30 s. Final extension was held at 72 ºC for two minutes, then brought down to 4 ºC and held for five minutes. Plates were then spun down at 1,000 rcf for one minute. We performed PCR cleanup using an AxyPrep PCR Clean-up Kit based on the manufacturer’s instructions, using the Viaflo multi-channel pipette. Samples were then indexed with the Nextera XT Index kit, following the manufacturer’s protocol, in 60 µl reaction volumes (9 µl sample, 6 µl each index, 39 µl GoTaq/Water mix).

To ensure that every sample was amplified to roughly the same DNA output, a round of qPCR was conducted using 18 µl of each sample with 2 µl of 10x SYBR Green. The qPCR protocol was: 72 ºC for 3 min, 95 ºC for 10 s, and 39 cycles of 95 ºC for 10 s, 55 ºC for 30 s, 72 ºC for 30 s. Final extension was 72 °C for 1 min and then 4 °C for 5 min. The mean Cq value (as described in the qPCR section) was calculated to determine the number of amplification cycles for the remaining 42 µL of sample. The amplification protocol itself was identical to the qPCR protocol (without plate reading and SYBR green labeling). Another round of PCR clean-up was conducted following the AxyPrep protocol described above. The resulting purified PCR products were then quantified using the Quant-iT dsDNA protocol and measured on a plate-reader with excitation and emission at 480 and 530 nm. Samples were then pooled at equimolar concentrations. The samples were quality-controlled by 10% native gel electrophoresis in TBE at 180 V for 40 min. The pool was then quantified again with Qubit and sent for sequencing on an Illumina MiSeq at the Stanford Functional Genomics Facility.

### HTS data analysis

FASTQ files were analyzed using custom python scripts. After demultiplexing in Basespace by Illumina indices, reads were filtered by quality, such that 90% of the base reads had to have a Phred score > 20. Filtered sequences were then assigned a sequence ID based on Smith-Waterman scoring. Striped Smith-Waterman scores for all sequences were calculated against the known aptamer or control sequences (**SI Table S1**). The best alignment score was then normalized by an optimal score (exact sequence match) and the sequence assignment was kept if the score was > 0.6. Otherwise, alignment was deemed poor and not assigned a sequence ID. The occurrence of each sequence per fastq file was then counted.

Each sample had a control sequence (CTRL.15 or CTRL.17) spiked in during HTS preparation prior to PCR. We use this to reduce intra-assay variance from experimental factors such as PCR bias, library prep, and pooling for sequencing. For each sample fastq file, the unique molecular ID count for the control sequence was counted based on the unique N15 region in the sequence (**SI Table S1**). All sequence counts (*apt*_*raw*_) were normalized by this control molecular barcode count (CTRL_*MBC*_):

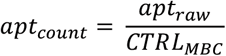

Finally, we calculated the total fraction of analyte bound:

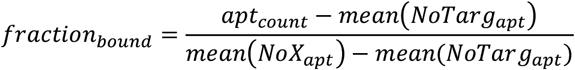

Where *mean(NoTarg_apt)* and *mean(NoX_apt)* are the mean counts of the aptamer in the no-target and no-crosslinking controls, respectively. The impact of this normalization by the control reads can be seen in **Figure S8**, compared to the normalized values in **Figure S7**.

### Curve-fitting

Binding curves for the dopamine dataset were modeled with the Langmuir isotherm:

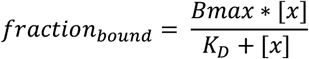

Where x is the analyte concentration, B_max_ is the maximum signal possible, and K_D_ is the dissociation constant. Since the kynurenine dataset has much more data per binding curve, we could fit with a slightly higher-order binding curve model without risking overfitting, and thus used the four-parameter logistic curve (4-PL):

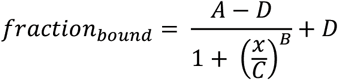

where fraction_bound_ is the normalized counts from sequencing, x is the analyte concentration, A is the minimum value possible with no analyte, B is the Hill coefficient, C is the point of inflection (K_D_), and D is the maximum value possible with infinite analyte. Curve-fitting was done with a custom python script, and all parameters were determined by Scipy’s optimize curve fit function, which uses non-linear least-squares to fit a function. We used the log residual for the loss function.

## Data Availability

The source and meta data generated in this study are provided on the Stanford Data Repository: https://purl.stanford.edu/rg880ws3662

## Code Availability

Code is available here: https://github.com/newmanst/xplex

## Acknowledgements

S.S.N acknowledges support from SGF (Stanford Graduate Fellowship in Science and Engineering) and the NSF Graduate Research Fellowship Program (GRFP). We also thank the Stanford Functional Genomics Facility and particularly Xuhuai Ji for their sequencing help.

## References

1. Darmanis, S. et al. Sensitive Plasma Protein Analysis by Microparticle-based Proximity Ligation Assays. Mol. Cell. Proteomics MCP 9, 327–335 (2010).

2. Lundberg, M., Eriksson, A., Tran, B., Assarsson, E. & Fredriksson, S. Homogeneous antibody-based proximity extension assays provide sensitive and specific detection of low-abundant proteins in human blood. Nucleic Acids Res. 39, e102 (2011).

3. Wu, Y., Tilley, R. D. & Gooding, J. J. Challenges and Solutions in Developing Ultrasensitive Biosensors. J. Am. Chem. Soc. 141, 1162–1170 (2019).

4. Seo, J. et al. PICASSO allows ultra-multiplexed fluorescence imaging of spatially overlapping proteins without reference spectra measurements. Nat. Commun. 13, 2475 (2022).

5. Nutiu, R. & Li, Y. Structure-Switching Signaling Aptamers. J. Am. Chem. Soc. 125, 4771–4778 (2003).

6. Yoshimura, Y. & Fujimoto, K. Ultrafast Reversible Photo-Cross-Linking Reaction: Toward in Situ DNA Manipulation. Org. Lett. 10, 3227–3230 (2008).

7. Sakamoto, T., Tanaka, Y. & Fujimoto, K. DNA Photo-Cross-Linking Using 3-Cyanovinylcarbazole Modified Oligonucleotide with Threoninol Linker. Org. Lett. 17, 936–939 (2015).

8. Nakatsuka, N. et al. Aptamer–field-effect transistors overcome Debye length limitations for small-molecule sensing. Science 362, 319–324 (2018).

9. Fornace, M. E. et al. NUPACK: Analysis and Design of Nucleic Acid Structures, Devices, and Systems. Preprint at 10.26434/chemrxiv-2022-xv98l (2022).

10. Yoshikawa, A. M., Wan, L., Zheng, L., Eisenstein, M. & Soh, H. T. A system for multiplexed selection of aptamers with exquisite specificity without counterselection. Proc. Natl. Acad. Sci. 119, e2119945119 (2022).

11. Newman, S. S. et al. Extending the dynamic range of biomarker quantification through molecular equalization. Nat. Commun. 14, 4192 (2023).

