## Supplemental Information for "A multiplexed and quantitative assay for small-molecule detection via photo-crosslinking of structure switching aptamers"

### Table S1 | DNA Sequences used in this work.

| Sequence Name | Sequence (5' -> 3') |
|---|---|
| Comp15_dop_glu | CAGCATGGGA/Xmod/GACG/3AmMO/ |
| Dop | GGCTCTCGGGACGACGCCAGTTTGAAGGTTCGTTCGCAGGTGTGGAGTGACGTCGTCCCATGCTG |
| Glu | GGCTCTCGGGACGACCGTGTGTGTTGCTCTGTAACAGTGTCCATTGTCGTCCCATGCTG |
| RP1 | GTCTCGTGGGCTCGGCAGCATGGGAC |
| FP1 | TCGTCGGCAGCGTCGGCTCTCGGGACGAC |
| PrimeTime_dop Probes | /56-FAM/TCCACACCT/Zen/GCGAACGAACC/3IABkFQ/ |
| CTRL.15 | GGCTCTCGGGACGACNNNNNNNNNNNNNNNGATAATTCCGTATTCCCTTTAATGATGTACGTCGTCCCTGAATTC |
| comp11 | CGTCCCGA/Xmod/AG/3AmMO |
| comp13 | GTCGTCCCGA/Xmod/AG/3AmMO |
| comp14 | TCGTCCCGA/Xmod/AGCC/3AmMO |
| 3HK-1 | GGCTCTCGGGACGACACGGGAAGCTTTAGGTTGAGCCATGTGCAGGTCGTCCCTGAATTC |
| XA-1 | GGCTCTCGGGACGACCGGAGGTCTCTTTACTTTTAACCAGGTGAGGTCGTCCCTGAATTC |
| KA-1 | GGCTCTCGGGACGACGATGGCGGTGTTTCTTTATTCGTAAATGGGGTCGTCCCTGAATTC |
| SK-1 | GGCTCTCGGGACGACGGTATTGCATCTTGGAATACAGCTTTGCTAGTCGTCCCTGAATTC |
| comp16 | GTCGTCCCGA/Xmod/AGCCT/3AmMO |
| comp17 | GTCGTCCCGA/Xmod/AGCCTG/3AmMO |
| 3HK.17 | CAGGCTCTCGGGACGACACGGGAAGCTTTAGGTTGAGCCATGTGCAGGTCGTCCCTGAATTC |
| XA.17 | CAGGCTCTCGGGACGACCGGAGGTCTCTTTACTTTTAACCAGGTGAGGTCGTCCCTGAATTC |
| KA.17 | CAGGCTCTCGGGACGACGATGGCGGTGTTTCTTTATTCGTAAATGGGGTCGTCCCTGAATTC |
| SK.17 | CAGGCTCTCGGGACGACGGTATTGCATCTTGGAATACAGCTTTGCTAGTCGTCCCTGAATTC |
| CTRL.17 | CAGGCTCTCGGGACGACNNNNNNNNNNNNNNNGATAATTCCGTATTCCCTTTAATGATGTACGTCGTCCCTGAATTC |
| FP17 | TCGTCGGCAGCGTCAGATGTGTATAAGAGACAGCAGGCTCTCGGGACGAC |
| RP2 | GTCTCGTGGGCTCGGAGATGTGTATAAGAGACAGGAATTCAGGGACGAC |

### Table S2 | 4-PL parameter fits and respective errors for dopamine and glucose aptamer and analyte pairs.

| Target | Aptamer | Min | Min Error | Hill | Hill Error | $K_D$ | $K_D$ Error | Max | Max Error |
|---|---|---|---|---|---|---|---|---|---|
| **dop** | **Dop** | **0.012** | **0.001** | **1.0** | **0.1** | **40 μM** | **6** | **0.634** | **0.034** |
| dop | Glu | 0.066 | 0.017 | 1.1 | 4.8 | 39 μM | 227 | 0.090 | 0.033 |
| glu | Dop | 0.012 | 0.001 | 1.1 | 1.9 | 239 mM | 673 | 0.026 | 0.015 |
| **glu** | **Glu** | **0.087** | **0.012** | **1.1** | **0.6** | **730 mM** | **1,637** | **2.182** | **2.395** |

**Table S3 | Tested mixtures of glucose and dopamine**

| Sample | [dopamine] (µM) | [glucose] (mM) |
|---|---|---|
| 1 | 5 | 5 |
| 2 | 5 | 50 |
| 3 | 5 | 500 |
| 4 | 50 | 5 |
| 5 | 50 | 50 |
| 6 | 50 | 500 |
| 7 | 500 | 5 |

**Table S4 | 4-PL parameter fits and respective errors for all kyn aptamer and analyte pairs**

| Target | Aptamer | Min | Min Error | Hill | Hill Error | $K_D$ (µM) | $K_D$ Error | Max | Max Error |
|---|---|---|---|---|---|---|---|---|---|
| **3HK** | **3HK-1** | **0.061** | **0.007** | **1.200** | **0.242** | **100** | **34** | **0.584** | **0.074** |
| 3HK | XA-1 | 0.022 | 0.018 | 1.143 | 2.090 | 37 | 79 | 0.103 | 0.061 |
| 3HK | KA-1 | 0.035 | 0.018 | 1.200 | 72.982 | 2,683 | 1,337,845 | 0.000 | 14.310 |
| 3HK | SK-1 | 0.024 | 0.009 | 1.200 | 0.806 | 149 | 199 | 0.292 | 0.158 |
| KA | 3HK-1 | 0.027 | 0.015 | 1.200 | 37.052 | 5,000 | 2,318,684 | 0.143 | 50.258 |
| KA | XA-1 | 0.500 | 252.684 | 1.200 | 12.121 | 0 | 32 | 0.016 | 0.005 |
| **KA** | **KA-1** | **0.078** | **0.008** | **0.900** | **0.264** | **325** | **294** | **0.572** | **0.177** |
| KA | SK-1 | 0.069 | 0.009 | 1.200 | 0.266 | 40 | 11 | 0.493 | 0.043 |
| XA | 3HK-1 | 0.035 | 0.006 | 0.900 | 2.321 | 665 | 5,883 | 0.083 | 0.184 |
| **XA** | **XA-1** | **0.042** | **0.008** | **1.183** | **0.218** | **48** | **14** | **0.611** | **0.064** |
| XA | KA-1 | 0.034 | 0.007 | 1.200 | 130.503 | 5,000 | 8,152,790 | 0.049 | 23.623 |
| XA | SK-1 | 0.032 | 0.005 | 1.200 | 0.281 | 131 | 60 | 0.541 | 0.102 |
| Kyn | 3HK-1 | 0.024 | 0.005 | 1.200 | 5.183 | 2,482 | 83,994 | 0.153 | 3.494 |
| Kyn | XA-1 | 0.016 | 0.004 | 1.142 | 1.003 | 660 | 1930 | 0.151 | 0.216 |
| Kyn | KA-1 | 0.021 | 0.006 | 1.200 | 18.377 | 5,000 | 1,155,800 | 0.123 | 22.206 |
| Kyn | SK-1 | 0.017 | 0.005 | 1.200 | 0.418 | 262 | 260 | 0.536 | 0.258 |

**Table S5 | 4-PL parameter fits and respective errors for Yoshikawa and Wan KA-1, 3HK-1, and XA-1 aptamers** We re-fit the original data provided by Yoshikawa and Wan with a 4-PL for direct comparison to the fit data we obtained with xPlex.

| Target | Aptamer | Min | Min Error | Hill | Hill Error | $K_D$ (µM) | $K_D$ Error | Max | Max Error |
|---|---|---|---|---|---|---|---|---|---|
| **XA** | **XA-1** | 9,434 | 98 | 0.9 | 0.1 | 375 | 144 | 22,214 | 2,072 |
| **3HK** | **3HK-1** | 14,729 | 97 | 1.2 | 0.1 | 7 | 1 | 22,108 | 461 |
| **KA** | **KA-1** | 16,754 | 102 | 1.2 | 0.1 | 48 | 4 | 25,950 | 230 |

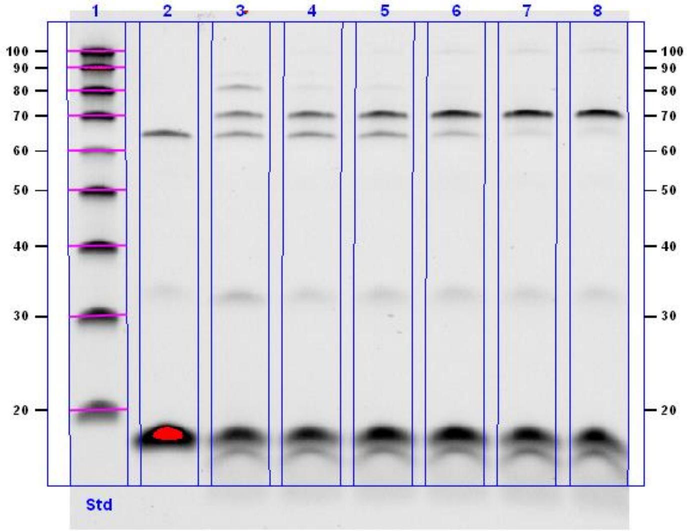


**Figure S1 | Dopamine binding to the dopamine aptamer in the presence of competitor.** Lanes 1–8 respectively show 20/100-nt ladder, non-crosslinked (NoX) control, and crosslinked aptamer-competitor complexes (50 nM aptamer and 2.5 µM competitor) challenged with 500, 100, 50, 10, 1, or 0 µM dopamine. Bands at < 20 nt represent the competitor strand. The slightly lower weight band seen in crosslinked samples is likely due to the formation of internal crosslinks within the competitor strand itself, which allow it to migrate faster than unlinked samples (note its absence in the NoX control, lane 2). Bands at ~70 and 65 nt are respectively crosslinked aptamers with competitor strand and unlinked aptamers. The high molecular weight band (> 80 nt) seen at high concentrations of dopamine is predicted to be arise from a second competitor strand that binds to the middle of the aptamer in addition to the normal binding region at the end of the aptamer, which is only predicted to become possible with weakened hairpin structure arising from high concentrations of dopamine.

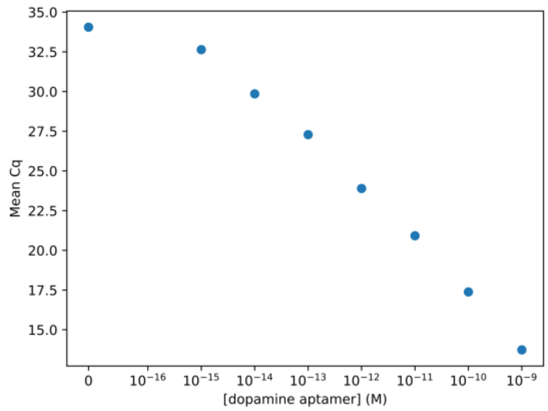


**Figure S2 | Dopamine PrimeTime calibration curve for qPCR efficiency calculation.** Increasing dopamine aptamer concentration decreases Cq value as determined by PrimeTime_dop probes.

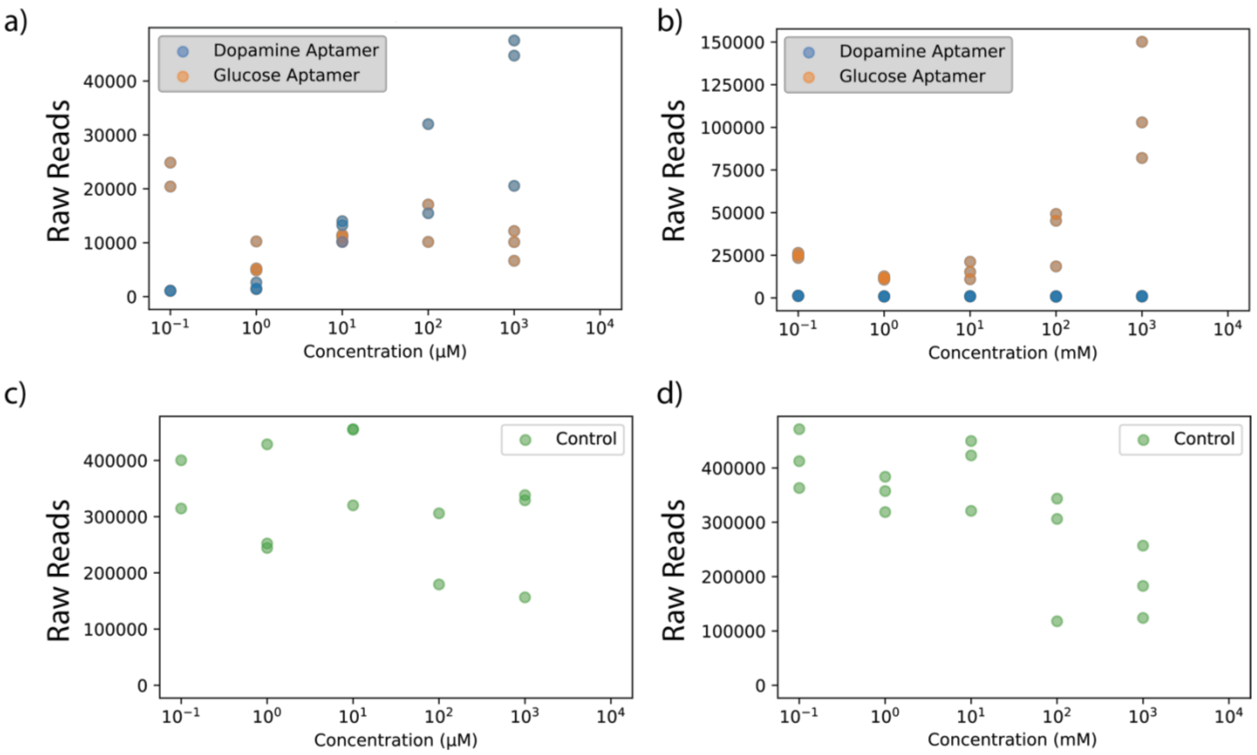


**Figure S3 | Non-normalized sequencing reads.** Binding curves for **a)** dopamine and **b)** glucose. **c-d)** Control sequence reads for respective dopamine and glucose samples.

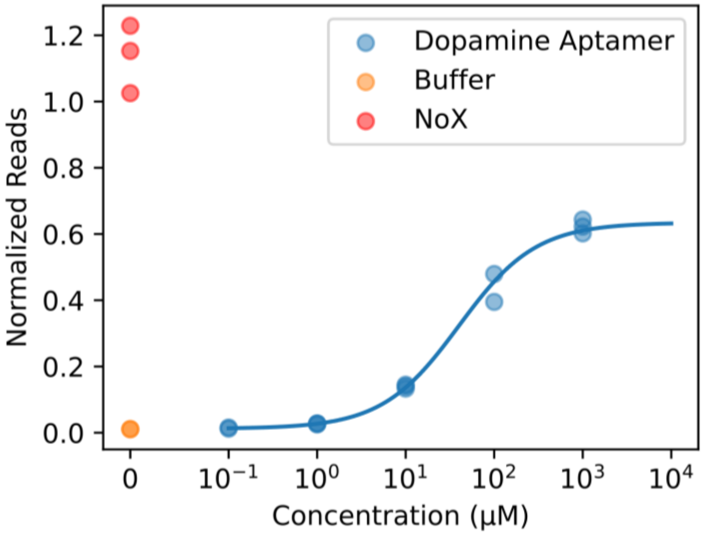


**Figure S4 | Dopamine binding curve with HTS readout with buffer (no analyte) and NoX (no crosslinking) controls.**

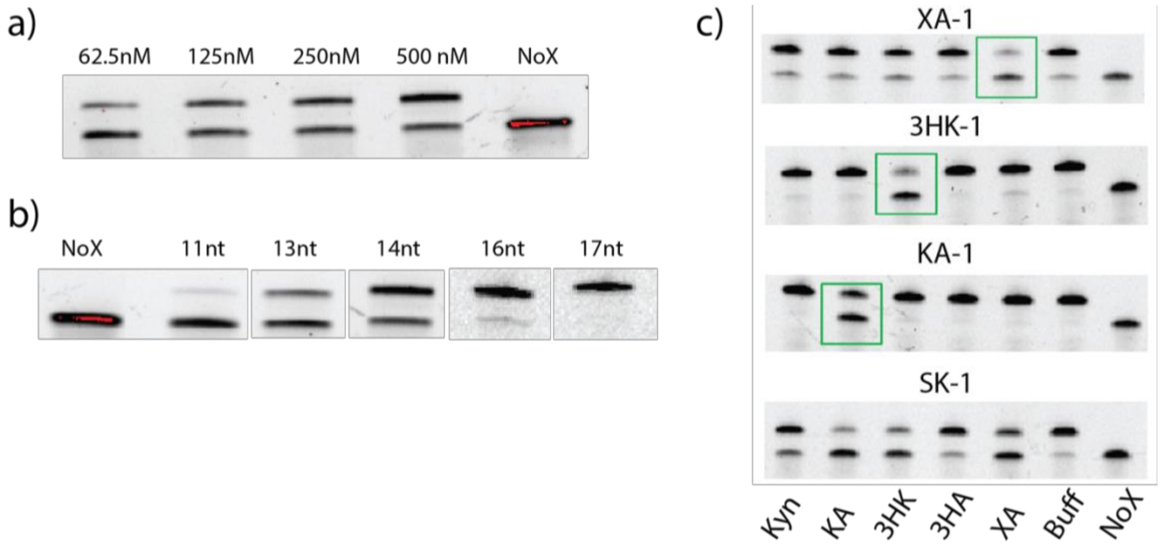


**Figure S5 | Gel characterization of competitor strand length and concentration for kynurenine pathway aptamers.** Gel-based characterization of **a)** competitor strand (Comp14) concentration and **b)** competitor strand length for XA-1 aptamer at 100 nM. Competitor strand concentration in the latter experiment was fixed at 500 nM. Red indicates signal saturation on the gel imager. **c)** Selectivity of XA-1, 3HK-1, KA-1, and SK-1 aptamers with 250 nM 16-nt competitor strand (Comp16) for 500 µM kyn, KA, 3HK, 3HA, or XA, plus buffer and non-crosslinked controls. Green box indicates the primary target for each aptamer. Full gel images and details are provided in **Figures S6–S8**. We saw a decrease in background from the unhybridized aptamer when we increased the concentration of the competitor strand for XA-1 (**a**) or increased the competitor strand length to further stabilize the competitor-aptamer duplex (**b**). We observed similar results when characterizing the 3HK-1, KA-1, and SK-1 aptamers (**Fig. S6–S8**).

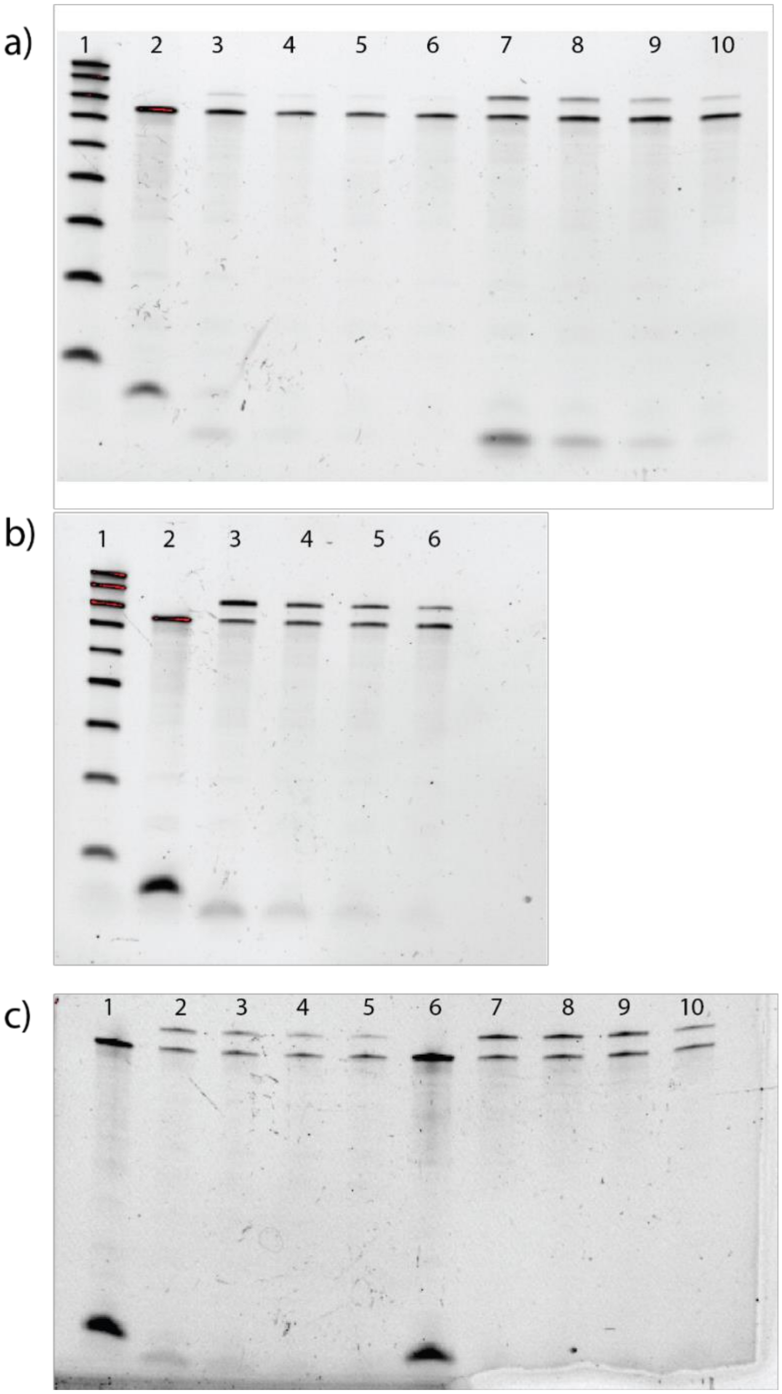


**Figure S6 | Characterizing competitor strands of varying lengths (11–14 nt).** Electrophoretic gels evaluating competitor strands measuring **a)** 11 and 13 nt or **b, c**) 14 nt in length. For **a**, lane 1 shows 20/100-nt ladder, lane 2 is the non-crosslinked control complex of aptamer and 14-nt competitor strand (comp14), lanes 3–6 respectively show XA-1 aptamer with 500, 250, 125, or 62.5 nM concentrations of 11-nt competitor strand (comp11), and lanes 7–10 show the same conditions with a 13-nt competitor (comp13). For **b**, lane 1 shows 20/100-nt ladder, lane 2 is the non-crosslinked control complex of aptamer and comp14, and lanes 3–6 respectively show XA-1 aptamer with 500, 250, 125, or 62.5 nM comp14. For **c**, lanes 1 and 6 show NoX control for 3HK-1 and KA-1, respectively, with comp14. Lanes 2–5 and 7–10 respectively show 3HK-1 and KA-1 with 1,000, 500, 250, and 125 nM comp14.

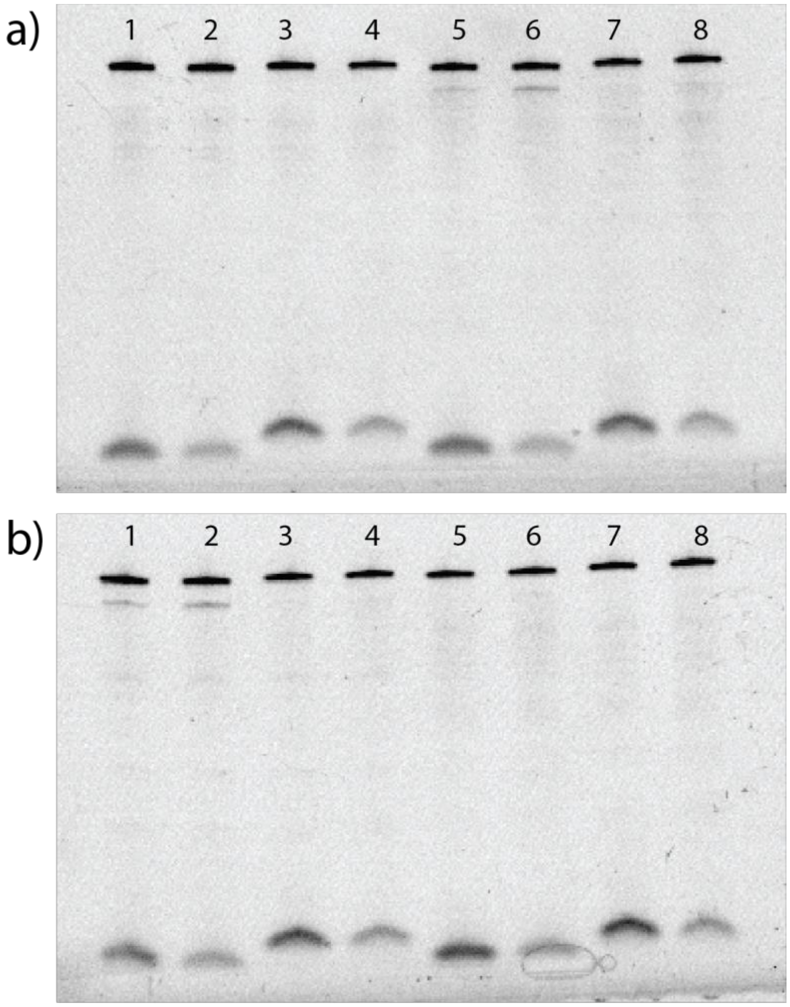


**Figure S7 | Characterizing competitor strands of varying lengths (16–17 nt). a**) Lanes 1 and 2 show aptamer XA-1 with 1,000 nM or 500 nM 16-nt competitor strand (comp16), and lanes 3 and 4 show the same conditions but with a 17-nt competitor strand (comp17). Lanes 5–8 show the same conditions, but with aptamer SK-1 instead of XA-1. **b**) Lanes show the same experimental conditions as in **a**, but with aptamer KA-1 in lanes 1–4 and aptamer SK-1 in lanes 5–8.

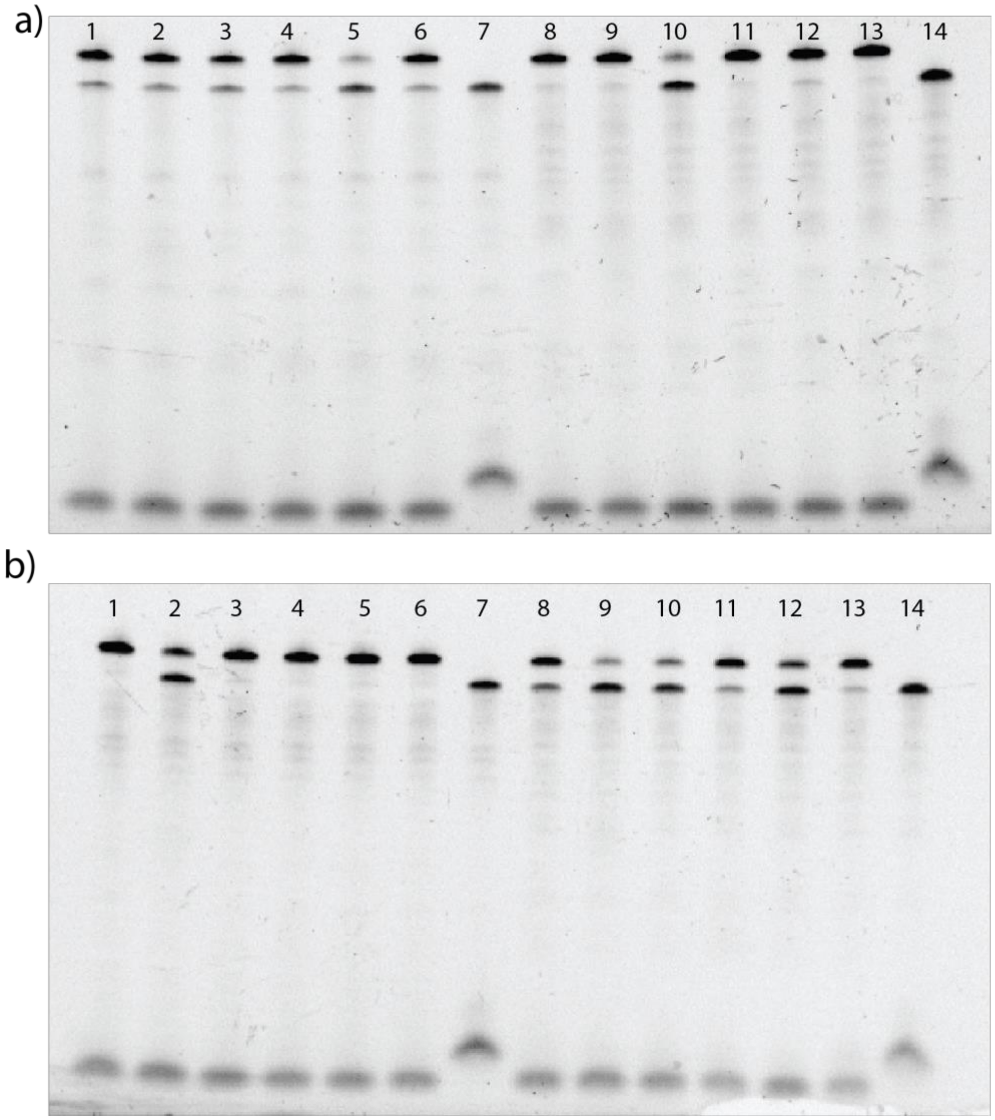


**Figure S8 | Characterizing the selectivity of kyn pathway aptamers.** Gel electrophesis analysis of aptamer response by **a**) XA-1 (lanes 1–7) and 3HK-1 (lanes 8–14) and **b)** KA-1 (lanes 1–7), and SK-1 (lanes 8–14) to kyn, XK, 3HK, 3HA, XA, buffer, and non-crosslinked controls, respectively. All experiments used 250 nM comp16 and 100 nM aptamer (XA_v2, 3HK_v2, KA_v2, and SK1_v2 in **SI Table S1**).

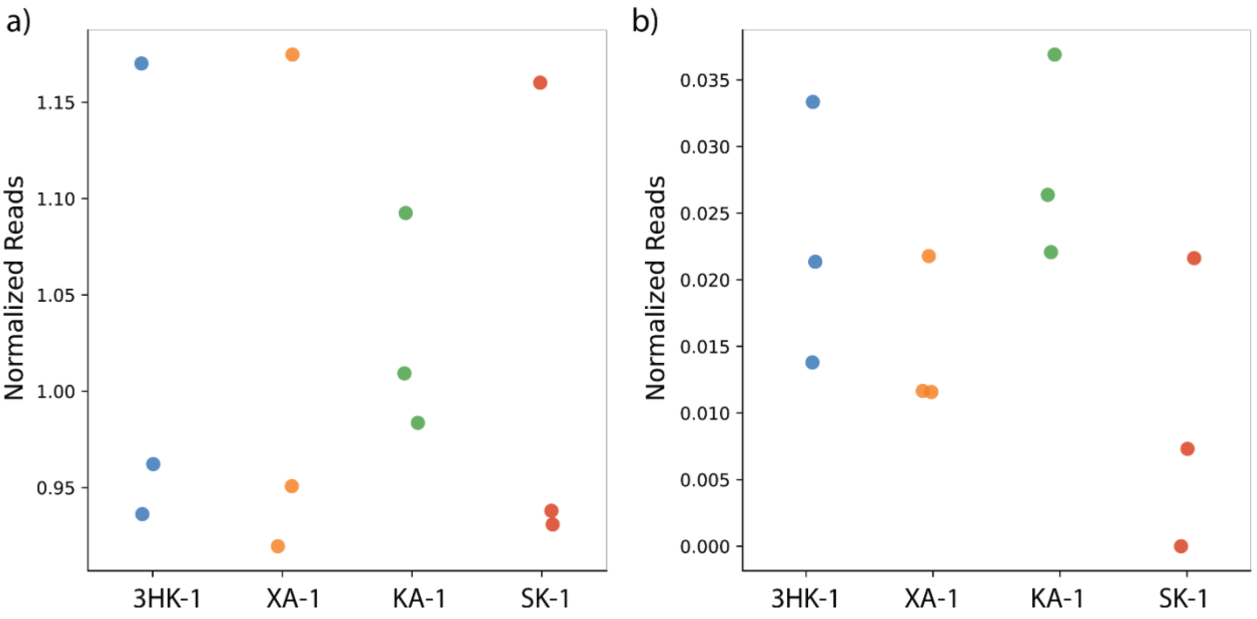


**Figure S9 | Kynurenine binding curves from HTS controls.** Normalized reads for each aptamer with no analytes **a**) without or **b**) with crosslinking.

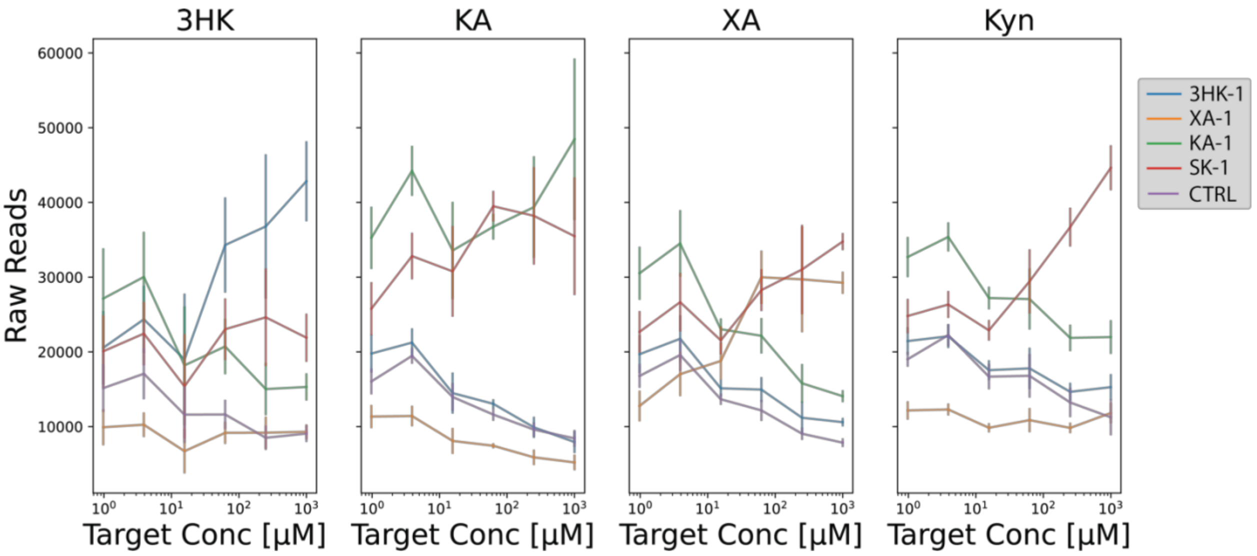


**Figure S10 | Non-normalized kynurenine binding curves for XA-1, 3HK-1, KA-1, and SK-1.** Raw signal readout before normalizing to the spiked control sequence (CTRL, purple).
